# Immune microenvironment subtypes and association with tumor cell mutations and antigen expression in follicular lymphoma

**DOI:** 10.1101/2021.04.27.441656

**Authors:** Guangchun Han, Qing Deng, Enyu Dai, Minghao Dang, John Ma, Haopeng Yang, Olga Kudryashova, Mark Meerson, Sergey Isaev, Nikita Kotlov, Krystle Nomie, Alexander Bagaev, Simrit Parmar, Fredrick Hagemeister, Sairah Ahmed, Swami Iyer, Filepe Samaniego, Raphael Steiner, Luis Fayad, Hun Lee, Nathan Fowler, Francisco Vega, Christopher R. Flowers, Paolo Strati, Jason R. Westin, Sattva S. Neelapu, Loretta J. Nastoupil, Linghua Wang, Michael R. Green

## Abstract

Follicular lymphoma (FL) is a B-cell lymphoma with a complex tumor microenvironment that is rich in non-malignant immune cells. We applied single-cell RNA-sequencing to characterize the diverse tumor and immune cell populations of FL and identified major phenotypic subsets of FL T-cells including a novel cytotoxic CD4 T-cell population. Their relative proportions of T-cells defined four major FL subtypes, characterized by differential representation or relative depletion of distinct T-cell subsets. By integrating exome sequencing, we observed that somatic mutations are associated with, but not definitive for, reduced antigen presentation on FL cells. In turn, expression of MHC class II genes by FL cells was associated with significant differences in the proportions and targetable immunophenotypic characteristics. This provides a classification framework of the FL microenvironment, their association with FL genotypes and antigen presentation, and informs different potential immunotherapeutic strategies based upon tumor cell MHC class II expression.

**Statement of significance:** We have characterized the FL-infiltrating T-cells, identified cytotoxic CD4 T-cells as an important component, showed that the abundance of these T-cell populations is associated with tumor-cell-intrinsic characteristics, and identified sets of targetable immune checkpoints on T-cells that differed between FLs with normal versus low antigen presentation.

## Introduction

Follicular lymphoma (FL) is an indolent lymphoma of germinal center B-cells that maintain follicle-like architecture and interact closely with T-cells and other immune cells. These immune interactions are critical to FL etiology^1^ and can be perturbed by somatic mutations that are frequent in FLs^2-4^. Understanding the immune tumor microenvironment (iTME) of FL and the interplay between perturbed immune interactions and distinct tumor-infiltrating T-cell (TINT) populations will be important for building precision immunotherapeutic approaches, but these concepts have yet to be comprehensively addressed using high-throughput approaches. Single cell RNA-sequencing (scRNA-seq) is a powerful and high-throughput approach that has revealed the deregulation of normal B-cell developmental programs and allowed for the characterization of targetable immune checkpoints on TINT cells^5,6^. However, these studies have been limited to a few patients and has not yet been used to investigate broader iTME profiles, or the relationship between somatic mutations, tumor B-cell expression profiles and changes in the iTME. Using scRNA-seq of FL lymph node biopsies, we characterized phenotypically distinct subsets of TINT cells, including a novel cytotoxic CD4 T-cell population, and validated in a large series that the composition of these T-cell subsets defines four distinct subtypes of iTME in FLs. By integrating exome sequencing and scRNA-sequencing data, we showed that somatic mutations in chromatin modifying genes can affect the expression of immune interaction genes encoding proteins such as major histocompatibility complex (MHC) class I and class II on tumor cells, which is in turn associated with changes in the frequencies and targetable immune profiles of T-cell subsets in FL tumors.

## Results

### Single cell RNA sequencing (scRNA-seq) of FL

We performed scRNA-seq of 20 FL and three reactive lymph nodes (RLN) using the 10X Chromium platform to profile the transcriptome in addition to T-cell receptor (TCR) and immunoglobulin (Ig) repertoires (Table S1). Additional marker genes were subjected to targeted sequencing by CapID, as previously described^7^. Each biopsy was analyzed fresh to retain cell types that are sensitive to cryopreservation, and included 11 previously untreated and nine relapsed FLs (median 1 line of prior therapy, range 1-6) that were grade 1-2 (n=14) or 3a (n=6). RLN (n=3) samples were included as controls. We sequenced a median of 6,138 (range; 635-11,070) cells per sample to a median of 57,933 (range; 49,833-324,873) reads per cells and detected a median of 1,115 (range; 447-2,979) genes per cell. After rigorous quality filtering, 137,147 cells were retained for subsequent analyses (Figure 1A). Unsupervised clustering analysis following batch effects correction identified six major cell lineages: B-cell, T-cell, monocyte/macrophage, follicular dendritic cell (fDC), plasmacytoid dendritic cell (pDC), and erythroid cell clusters, as determined by cluster marker genes (**Figure 1B-C**; Table S2).

**Figure 1:**
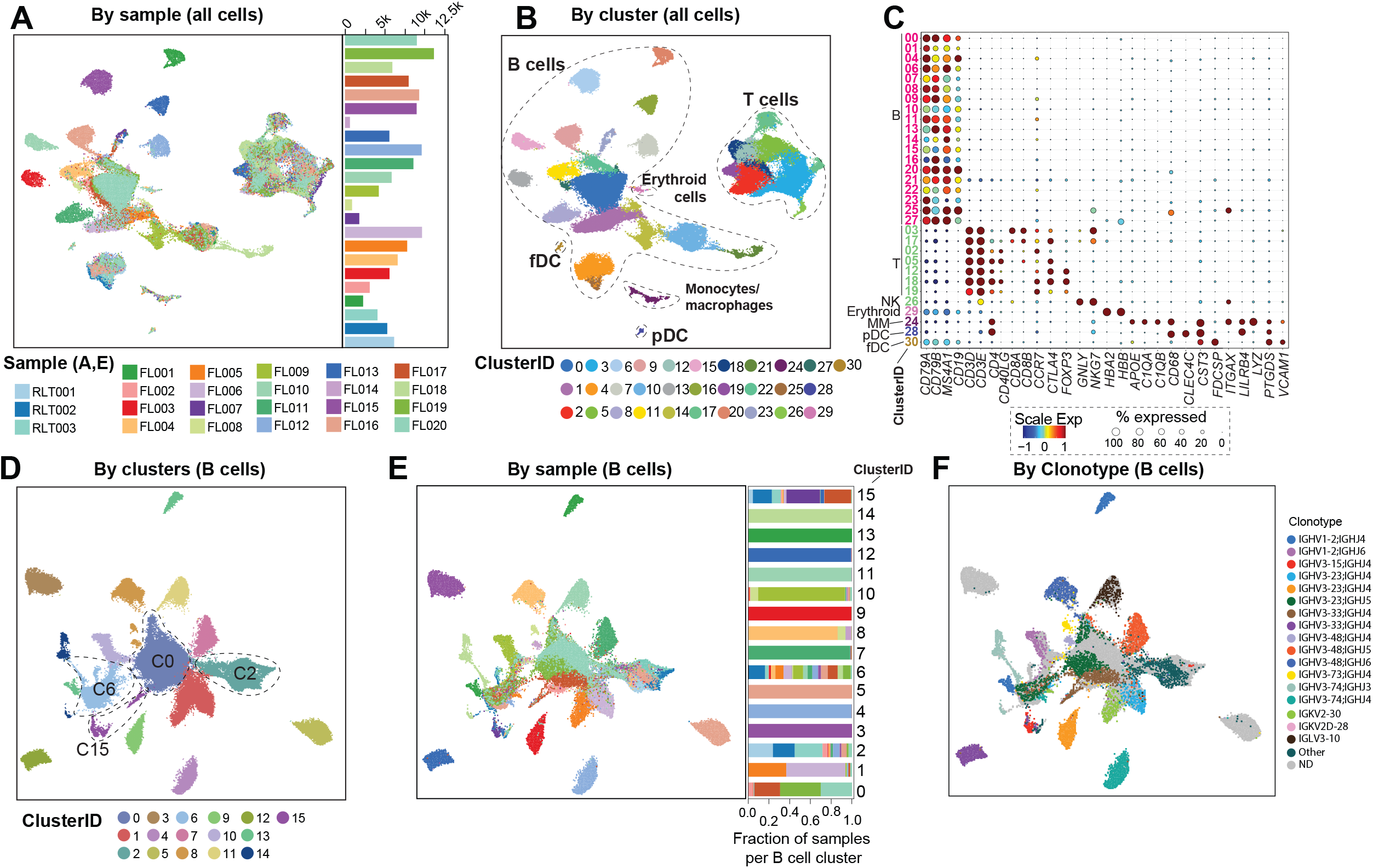
Overview of major cell types and clusters from single cell RNA sequencing of 20 FL tumors. **A-B)** UMAP plots show 137,147 cells from 20 FL tumors and 3 RLT controls by sample ID (A) and cluster ID (B). Major cell types are annotated in B. **C)** Bubble plot of cell lineage marker genes are shown for B-cell, T-cell, natural killer cell (NK), erythroid, monocyte/macrophage (MM), plasmacytoid dendritic cell (pDC) and follicular dendritic cell (fDC) clusters. **D-F)** UMAP plots show re-clustering of 99,610 B-cells by cluster ID (D), sample ID (E), and immunoglobulin clonotype (F). Among B-cell clusters, we identified those corresponding to non-malignant B-cells (C2), proliferating cells (C6), plasma cells (C15). A malignant B-cell cluster bearing cells from multiple samples was identified (C0). The contribution of each sample to each cluster is shown in the bar graph in E, with many clusters consisting of tumor B-cells from a single sample as determined by immunoglobulin clonotype (F) or patterns of inferred copy number variation (Figure S1).

B-cells were re-clustered (**Fig. 1D-E**) and cells defined as either tumor or non-malignant by the presence/absence of a clonal immunoglobulin sequence (**Fig. 1F**) or DNA copy number alterations (Fig. S1). Clusters of non-malignant B-cells (C2), plasma cells (C15) and proliferating B-cells (C6) included non-malignant cells from both FL and RLN samples (**Fig. 1D-E**). A central cluster (C0) was also found to contain cells from multiple samples, but consisted exclusively of clonal malignant B-cells from FLs, suggesting that tumor cells from a subset of cases have shared transcriptional characteristics. These FLs consisted of both low and high-grade tumors, but 74% of cells originated from treatment-naïve tumors (Table S1) suggesting that tumor B-cells from relapsed FLs have a greater inter-sample divergence in transcriptional profiles compared to treatment-naïve FLs.

### Tumor infiltrating T-cell composition defines iTME subtypes of FL

T-cells comprised of a median of 87.6% (range 73.8% to 98.9%) of the non-malignant cells within the iTME (**Fig. 2A**). We further characterized phenotypically distinct subsets of CD4 and CD8 T-cells by subclustering analysis (Fig. S2; Table S3). Clusters of CD8 T-cells included naïve (*CCR7, SELL*, and *IL7R*), effector (granzymes *GZMA/B/K* and *PRF1*) and exhausted (CD8_Exh_, high expression of inhibitory immune checkpoint genes such as *TIGIT* and *LAG3*, and a high exhaustion score) subsets (**Fig. 2B-C**). Trajectory analysis showed that these represent a functional continuum from naïve through to exhausted states (Fig. S3). Subclustering analysis of CD4 T-cells identified four transcriptome states (**Fig. 2D**), including naïve (high expression of *CCR7, SELL* and *IL7R*), T-regulatory (T_REG_; high expression of *FOXP3, CTLA4, IL2RA*), T follicular helper (T_FH_; high expression of *PDCD1, TOX, TOX2, CXCR5* and *CD40LG*), and cytotoxic CD4 T-cells (CD4_CTL_; high expression of *GZMA*/K, *NKG7*, and *EOMES*), all of which were detected in both FL and RLN samples. While naïve, T_REG_ and T_FH_ cells are well-described components of FL^1^, there are no prior reports of CD4_CTL_ cells in FL or any other germinal center derived lymphoma. CD4_CTL_ cells express *CD4* but not *CD8A/B*, have a high cytotoxicity score with *GZMK* expression detectable in 89.6% of cells, and high expression of the *EOMES* transcription factor that is implicated in CD4_CTL_ development^8^. In addition, CD4_CTL_ cells bear some similarities to T_FH_ cells, including high expression of *CXCL13* and *PDCD1*, and are most closely related to T_FH_ cells by trajectory analysis (Fig. S3). A high fraction of CD4_CTL_ expressed co-inhibitory receptors (*LAG3, CTLA4, HAVCR2*; Table S3) that are potentially targetable. Thus, our scRNA-seq analysis revealed a novel cytotoxic CD4 T-cells component of the lymphoid and FL iTME that requires further functional exploration.

**Figure 2:**
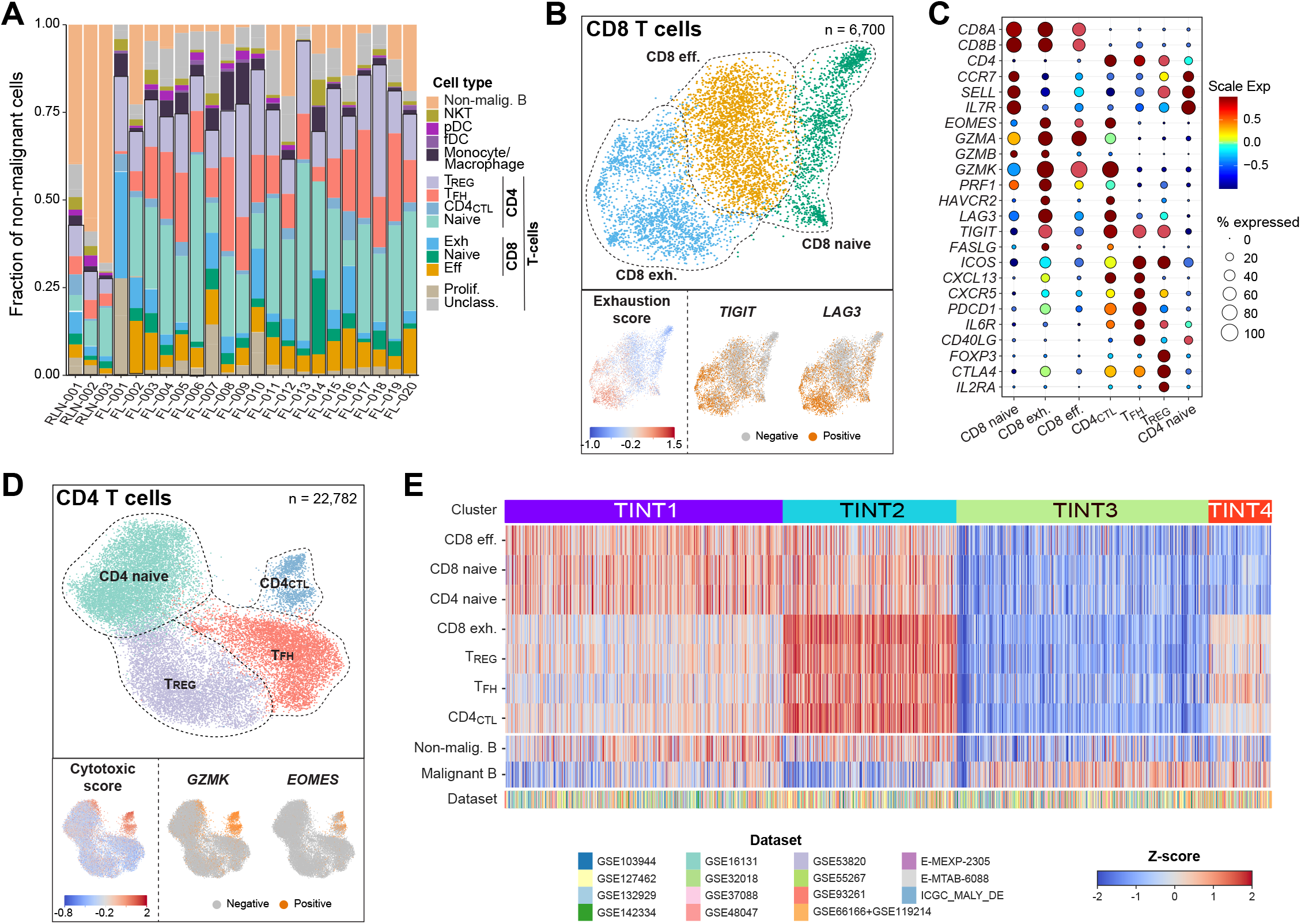
Tumor infiltrating T-cell populations in follicular lymphoma. **A)** A bar graph shows the frequency of non-malignant immune cell populations within FL, with the majority of cells belonging to the T-cell lineages. **B)** UMAP plots from re-clustering of 6,700 CD8 T-cells shows 3 major populations aligning with naïve, effector (eff) and exhausted (exh) states. Single cell GSVA of a CD8 T-cell exhaustion signature shows the highest expression in the CD8_Exh_ cluster, which is also characterized by high expression of *TIGIT* and *LAG3*. **C)** A bubble plot shows the proportion of cells of CD8 and CD4 T-cell clusters expressing known phenotypic marker genes (size of circles) and their average expression levels (color of circles). D) UMAP plots from re-clustering of 22,782 CD4 T-cells shows 4 major subpopulations aligning with naïve, regulatory (T_REG_), T follicular helper (T_FH_) and CD4 cytotoxic (CD4_CTL_) states. Single cell GSVA of a cytotoxic score including immune effector molecules shows high expression in the CD4_CTL_ cluster, which is also characterized by high expression of *GZMK* and *EOMES*. **E)** A heatmap shows the relative proportions of CD8 and CD4 tumor infiltrating T-cell (TINT) populations calculated by deconvolution from publicly available bulk gene expression microarray or RNA-sequencing datasets (n=1,269 FL tumors from 15 datasets, see Supplementary Methods). Unsupervised clustering identified 4 characteristics patterns (TINT1-4) with different relative abundance of tumor-infiltrating T-cell populations.

The abundance of functionally distinct tumor infiltrating T-cell (TINT) populations that we characterized by scRNA-seq were highly variable across patients (**Fig. 2A**). We therefore assessed their representation across an external validation set of bulk gene expression profiling (GEP) from 1,269 FLs compiled from 15 datasets. Signatures derived from our scRNA-seq data were validated in publicly available scRNA-seq data (Fig. S4), then used to infer the abundance of each cell type by single cell gene set enrichment analysis (ssGSEA) followed by clustering the inferred frequencies to define sets of tumors with similar TINT profiles (**Fig. 2E**), as previously described^9^. This revealed four distinct subtypes of iTME in primary human FL based on the relative abundance of TINT cells: (TINT1) high in CD8 effector, CD8 naïve and CD4 naïve; (TINT2) high in CD8_Exh_, T_REG_, T_FH_ and CD4_CTL_; (TINT3) high in malignant B-cells and depleted of T-cell subsets; (TINT4) high in malignant B-cells and depletion of CD8 effector, CD8 naïve and CD4 naïve. The landscape of TINT as defined by scRNA-seq cell composition and measured in bulk GEP data therefore defines four distinct subsets of iTME in primary human FL.

### Multiple mechanisms of MHC class II loss on FL tumor B cells

Mutations in chromatin modifying genes (CMGs) are a hallmark of FL^10^, and affect the expression of genes in tumor B-cells through epigenetic dysregulation. The most frequently mutated CMGs (*KMT2D, CREBBP* and *EZH2*) have each been implicated in deregulating interactions between tumor cells and T-cells^3,4,11^, leading us to hypothesize that these mutations may underlie tumor-cell-intrinsic gene expression changes that drive differential TINT profiles. Using whole exome sequencing of tumors with available DNA (n=19; Table S5; **Fig. 3A**), we applied single cell differential gene expression profiling to identify genes that were significantly altered in association with these mutations (**Fig. 3B;** Tables S6). Collectively, the union of genes with significantly reduced expression (FDR q-value<0.05, fold change>1.2; n=355; Table S6) in association with one or more of these mutations was significantly enriched for genes involved in immune cell interactions (p = 1.4×10^−7^) including those with a role in antigen processing and presentation (p = 2.2×10^−29^), confirming that these mutations alter genes involved in immune cell interactions (**Fig**.**3C**). In line with prior reports, *CREBBP* and *EZH2* mutations were both associated with reduced expression of multiple genes involved in antigen presentation through the MHC molecule^3,11^, which present antigens that are recognized by T-cell receptors and therefore affect T-cell activation. Mutations of *CREBBP* co-occurred with *EZH2* mutations in three out of four cases and were predominantly associated with lower MHC class II (MHCII) expression (**Fig. 3D**), while *EZH2* mutations were selectively associated with lower MHC class I (MHCI) expression. *KMT2D* mutations were also associated with reduced expression of a subset of MHCI genes, and co-occurred with *EZH2* mutations in three out of four tumors. Using non-malignant B-cells from RLNs as reference to define normal *MHCI* and *MHCII* expression levels, we observed that loss of MHCI and/or MHCII was not restricted to *EZH2* and/or *CREBBP* mutant tumors (**Fig. 3E**). Specifically, MHCII loss was most prevalent and observed in 58% (11/19) of tumors, but 27% (3/11) of MHCII-low tumors lacked *CREBBP* or *EZH2* mutations. Further, one *CREBBP* mutant tumor did not show MHCII loss at mRNA level. CMG mutations in FL are therefore associated with perturbed expression of immune interaction genes on tumor B-cells, but additional mechanisms exist for MHCI and MHCII loss that are likely to have an equal impact on tumor infiltrating immune cells via deregulation of immune synapse formation.

**Figure 3:**
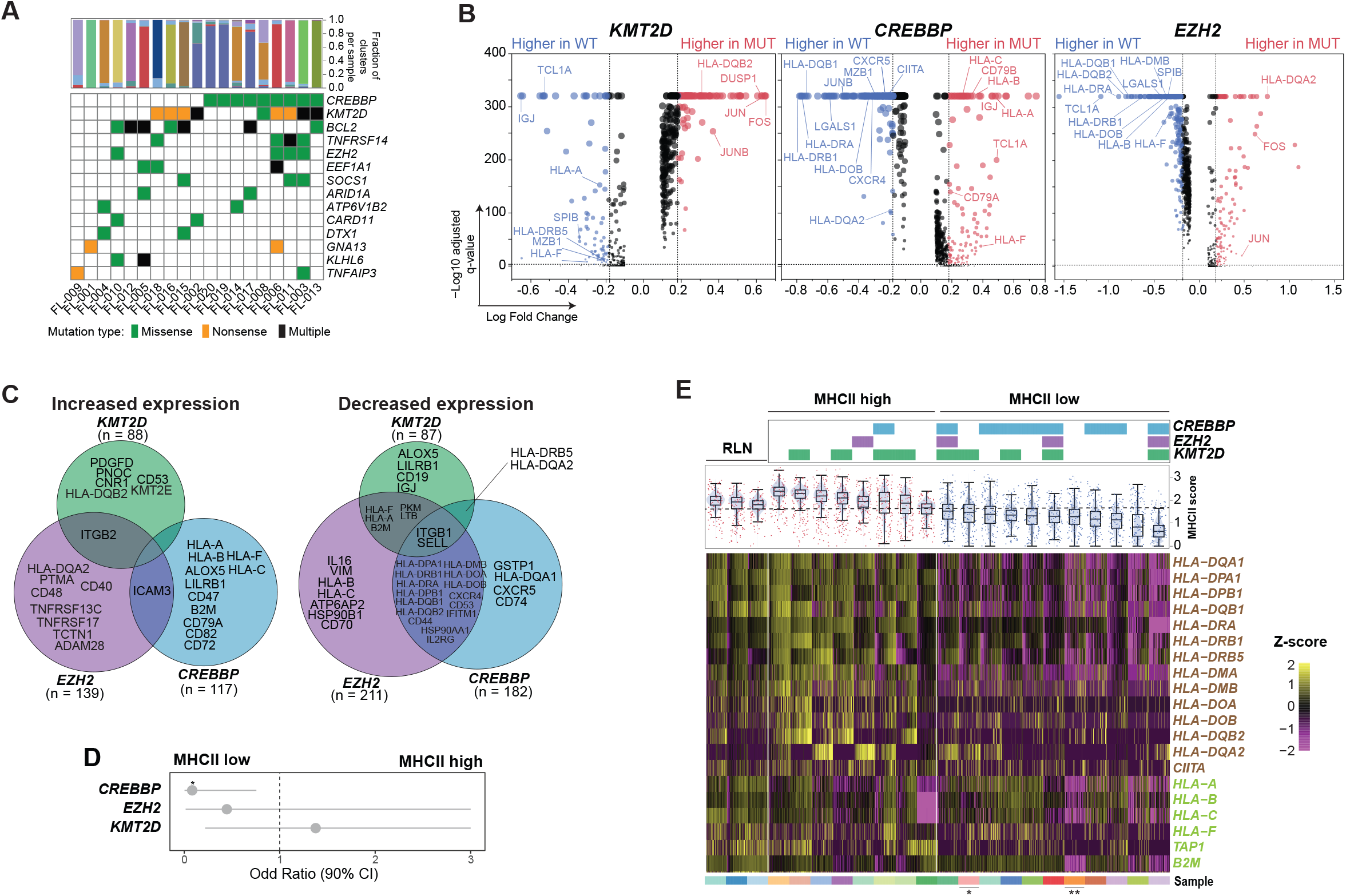
Effect of somatic mutations on tumor B-cell expression profiles. **A)** An oncoplot shows recurrently mutated genes in the 19 FL tumors with available DNA. B) Volcano plots displaying differentially expressed genes between tumor B-cells from *KMT2D* (left), *CREBBP* (middle) or *EZH2* (right) wild-type and mutant tumors. Examples are annotated and the full list provided in Table S6. **C)** Venn diagrams display the overlap of genes with increased (left) or decreased (right) expression associated with each mutation, with genes encoding cell surface proteins annotated. **D)** Odds ratio (+/- 95% CI) are shown for association between individual mutations and MHCII status (two-tailed Fisher’s exact test p=0.028). **E)** The expression of MHCII (brown, above) and MHCI genes (green, below) are shown for individual tumor B-cells from each tumor with available mutation data. Mutations of *CREBBP, EZH2* and *KMT2D* are annotated at the top. Sample IDs are colored according to Figure 1A and tumors with additional mutations in \**CIITA* and \*\**B2M* that may also affect MHCII and MHCI expression, respectively, are annotated by asterisk.

### Frequencies and targetable features of TINT are associated with tumor B-cell MHCII expression

Having observed different patterns of TINT in FL, and mutation-associated changes in MHCI and MHCII expression on tumor B-cells, we next evaluated whether these features were associated. Tumor MHCII loss was more significantly associated with TINT frequencies than somatic mutations of *CREBBP, EZH2* or *KMT2D* (Table S7), and was more frequent than MHCI loss, so we focused on this feature. MHCII-low tumors had significantly reduced levels of CD8_Exh_ and CD4_CTL_ (**Fig. 4A**) – features of the TINT2 microenvironment subtype (**Fig. 2E**). Despite a relatively modest sample size, we observed both a quantitative and qualitative relationship between MHCII expression/status and the frequencies of these TINT subsets (**Fig. 4B-D**). In mantle cell lymphoma, tumor cell immunopeptidome profiling revealed presentation of tumor idiotype peptides in MHCII that were recognized by CD4_CTL_ in the peripheral blood^12^. We therefore reasoned that loss of MHCII may be selectively acquired in cells that have accumulated immunogenic mutations in their idiotype sequences, and thus may be restricted to immunogenic clades of the immunoglobulin hierarchy. By evaluating paired single cell BCR sequencing data, we found anecdotal evidence of this in three FL tumors (Fig. S5), but this trend was not widespread in this cohort.

**Figure 4:**
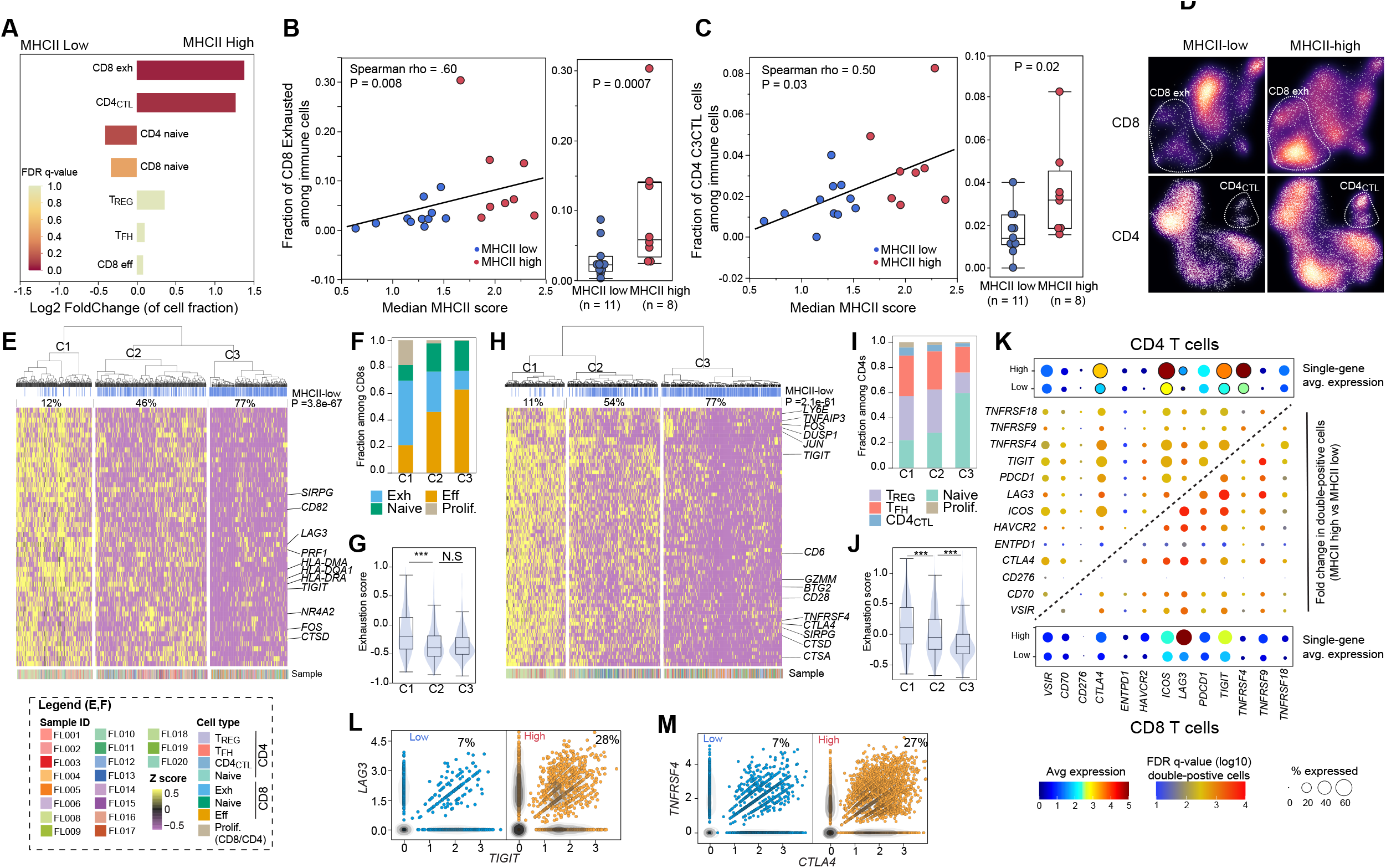
Association between tumor MHCII status and tumor infiltrating T-cell populations. **A)** A bar graph shows the fold change of CD8 and CD4 T-cell populations between MHCII low and MHCII high tumors, colored by Fisher Exact FDR q-value. The CD8 exhausted (exh; fold-change 3.43, Fisher exact q=0.05) and CD4_CTL_ (fold-change 2.11; Fisher exact q=0.07) populations are significantly higher in MHCII high tumors compared to MHCII low tumors. **B-C)** Scatter plots and bar plots show the quantitative and qualitative association between CD8 exhausted (B) and CD4_CTL_ (C) populations and either the expression or status of MHCII on tumor B-cells, respectively. **D)** UMAP density plots show the relative representation of CD8 exhausted (exh; above) and CD4_CTL_ (below) populations between MHCII low (left) or MHCII high (right) tumors. **E)** Differentially expressed genes (DEG) between CD8 T-cells from MHCII low vs MHCII high tumors were subjected to unsupervised hierarchical clustering identifying 3 clusters with significantly different proportions of cells from MHCII low/high tumors (top track, P=3.8×10^−67^). **F)** A bar graph of cell states in each DEG cluster from E, which shows higher fractions of CD8 exhausted (exh) cells in C1 and a higher fraction of CD8 effector cells (eff) cells in C3. **G)** GSVA showed higher expression of exhaustion signature genes in cells within C1 compared to either C2 or C3. **H**) Differentially expressed genes (DEG) between CD4 T-cells from MHCII low vs MHCII high tumors were subjected to unsupervised hierarchical clustering identifying 3 clusters with significantly different proportions of cells from MHCII low/high tumors (top track, P=2.1×10^−61^). **I)** A bar graph of cell states in each DEG cluster from H, which shows higher fractions of CD4_CTL,_T_FH_ and T_REG_ cells in C1 and a higher fraction of CD4 naïve cell in C3. **J)** GSVA showed higher expression of exhaustion signature genes in cells within C1 compared to either C2 or C3. **K)** A bubble plot shows the average expression of immune-modulatory genes on CD4 (above) or CD8 (below) T-cells in MHCII low or high tumors. The center grid shows the fold-change and significance of pairs of immune-modulatory genes between MHCII low and high tumors for CD4 (top left) and CD8 (bottom right) T-cells. **L-M)** Scatter plots show the co-expression of significant pairs of targetable immune modulatory genes in CD8 (L, *LAG3* and *TIGIT*) and CD4 (M, *TNFRSF4* and *CTLA4*) T-cells from MHCII low (left) or MHCII high (right) tumors.

In addition to changes in the frequencies of CD4 T-cells, we explored differences in gene expression of tumor infiltrating CD4 and CD8 T-cells using single cell differential gene expression analysis (Table S8-S9; **Fig. 4E-J**). Cells were clustered within the space of the differentially expressed genes (DEGs), which revealed three clusters for both CD4 and CD8 T-cells that had significantly different representation of cells from MHCII-high vs MHCII-low tumors (**Fig. 4E**, CD4, p=3.8×10^−67^; **Fig. 4H**, CD8, p=2.1×10^−61^). The DEGs includes markers of activation, transcription factors and multiple targetable cell surface immune checkpoint molecules. C1 clusters which have the lowest frequency of cells from MHCII-low tumors expressed the highest level of these genes, and C3 clusters which have the greatest frequency of cells from MHCII-low tumors express low levels of these genes. This is suggestive of higher levels of T-cell activation and exhaustion in tumors that have retained MHCII expression, as supported by GSVA analysis of a previously described exhaustion score (**Fig. 4G & J**), and in line with prior associations between MHCII expression and superior response to immune checkpoint blockade^13^. We therefore aimed to assess the most dynamic pairs of immune checkpoints that may serve as therapeutic targets in FL tumors with high MHCII (**Fig. 4K**). Within the CD8 T-cell compartment, the most significant change was increased frequencies of *LAG3* and *TIGIT* dual-expressing cells (fold-change = 4.3; FDR q-value = 1.6×10^−3^; **Fig. 4K & L**), which have yet to be explored as combination therapeutic targets in lymphoma. The most significant change in the CD4 T-cell compartment was the increased prevalence of *TNFRSF4* (aka. *OX40*) and *CTLA4* dual-expressing CD4 T-cells (fold-change = 3.9; FDR q-value = 0.01; **Fig. 4K & M**), combined targeting of which has been shown to be highly efficacious in preclinical models of lymphoma^14^. Thus, tumor cell MHCII expression correlates with the frequency and targetable immune profile of TINT cells in FL, highlighting subsets of FL that are likely to have differential responses to specific immune checkpoint blockade.

## Discussion

Follicular lymphoma is an indolent disease, with some patients having equivalent overall survival to age-matched controls^15,16^. Decreasing the use of cytotoxic chemotherapy in the treatment of FL is therefore a priority. The iTME of FL is a complex ecosystem that includes large numbers of T-cells that provide survival signals that are integral to disease etiology, offering an attractive opportunity for immunotherapeutics that target critical nexuses. However, single agent checkpoint blockers such as anti-PD1/PD-L1 are largely ineffective in FL^17^. Understanding the characteristics of the FL iTME and how it is modulated by tumor-cell-intrinsic characteristics is therefore an important step towards the rational design of combination immunotherapeutic strategies that may have increased efficacy.

The large number of cells that we sequenced afforded us the power to identify functionally distinct subsets of T-cells. Among these was a subset of CD4_CTL_ that have not been previously appreciated as a component of the FL iTME, and have been infrequently described in other cancers such as bladder cancer^18^ and in the peripheral blood of mantle cell lymphoma patients^12^. In the latter, these cells were shown to recognize tumor idiotype peptides presented in MHCII. However, we did not find strong evidence in support of this in FL. CD4_CTL_ play an important role in antiviral immune responses^8^, and their development in this context has been shown to be mediated by the transcription factors T-bet or EOMES^8^. Consistent with this, we observed high expression of *EOMES* in the CD4_CTL_ that we defined. Interestingly, CD4_CTL_ were also detected within RLN samples suggesting that these cells may be a normal component of the lymphoid microenvironment. However, there were significant differences in gene expression between CD4_CTL_ from FLs compared to RLN such as the downregulation of costimulatory receptors and IL6 signaling genes that are suggestive of dysfunction in FL. In addition, we identified multiple potential therapeutic targets on CD4_CTL_, including exhaustion markers *CTLA4, LAG3* and *HAVCR2* (aka. *TIM-3*). Future studies are needed to characterize the role of CD4_CTL_ in normal and malignant lymphoid tissues, and whether these cells can be targeted to induce anti-lymphoma immunity.

Loss of antigen presentation is common in FL and has been linked to recurrent mutations in *CREBBP* and *EZH2*^3,11^. We confirmed this association but also identified multiple cases of FL with mutation-independent loss of antigen presentation and showed that the antigen presentation status is more significantly associated with TINT characteristics than somatic mutations. Specifically, we observed an association between normal MHCII expression on tumor B-cells and higher frequencies of CD4_CTL_ and CD8_Exh_ T-cells. The high expression of exhaustion markers on both CD4_CTL_ and CD8_Exh_ suggests that FL tumors with normal MHCII expression may have an inflammatory microenvironment that promotes adaptive immune suppression and T-cell exhaustion. In other cancers, ‘warm’ microenvironments such as this show greater response to immune checkpoint blockade^19^. We explored potential therapeutic targets on the T-cells from FL tumors with retained MHCII expression and identified *LAG3*+*TIGIT* and *CTLA4*+*TNFRSF4* as potential combination immunotherapy targets for CD8 and CD4 T-cells, respectively. Our data also suggest that tumors with MHCII loss may have ‘cold’ microenvironments and be less responsive to immune checkpoint blockade. Therefore, tumor cell MHCII expression status should be prospectively explored as a potential biomarker for selection of, and response to, immune checkpoint therapies in FL.

CD19 chimeric antigen receptor (CAR) T-cell therapy is highly efficacious in relapsed/refractory FL and has recently been FDA approved in this setting. Responses are likely to be impacted by the tumor microenvironment characteristics of FL, but these characteristics have not been thoroughly explored in a large series of tumors. We therefore leveraged our signatures from scRNA-seq data to explore the relative representation of T-cell subsets in a large number of tumors using bulk GEP data. This identified four major subtypes of FLs characterized by different patterns of TINT cells, including ‘warm’ (TINT1, TINT2), ‘cold’ (TINT3) and intermediate (TINT4) subtypes, consistent with prior observations using NanoString GEP^20^. Mutation data were not available for these tumors to evaluate the relationship between tumor microenvironment subtype and mutations of *CREBBP* or *EZH2*, and tumor MHCII status cannot be predicted due to highly variable frequencies of tumor infiltrating T-cells and other antigen presenting cells. This will therefore require prospective validation using orthogonal approaches. However, consistent with our scRNA-seq data, T-cell subsets that express high levels of exhaustion markers (CD4_CTL_ and CD8_Exh_) were correlated in their relative representation across these microenvironment subtypes. We therefore suggest that evaluation of these tumor infiltrating T-cell subtypes may be important to prospectively evaluate in FL patients being treated with CD19 CAR T-cells and other cellular therapies or immunotherapies.

In conclusion, the FL tumor microenvironment is highly variable across patients and influenced by tumor-cell-intrinsic characteristics such as somatic mutations and antigen presentation status. The characteristics of tumor infiltrating T-cells allow for data-driven selection of combination immunotherapy targets, and highlight ‘warm’ and ‘cold’ microenvironments that are important to prospectively consider as potential determinants of immunotherapeutic and cellular therapy responses in FL patients.

## Methods

For detailed methods, please refer to the supplementary information. FL and RLN biopsies were obtained following informed consent under protocols approved by the Institutional Review Board of MD Anderson Cancer Center (Protocols 2005-0656 and PA19-0420). Tissues were processed fresh by physical disaggregation through a metal screen followed by a 40µM filter and loaded onto a 10X Chromium with 5’GEX chemistry to obtain a goal of 10,000 cells per sample. Transcriptome, BCR and TCR libraries were prepared and sequenced according to the manufacturer’s protocol. CapID hybrid-capture sequencing of transcriptome libraries was performed as previously described^7^. Single cell RNA-sequencing analysis was performed following quality filtering and batch correction using Seurat^21^. Genomic DNA from residual cells and interrogated by whole exome sequencing using Nimblegen SeqCap Exome v3. Somatic mutations were identified and annotated as previously described^22^.

## Acknowledgements

This work was supported by R01 CA201380 (MRG), the MD Anderson Cancer Center Support Grant (P30 CA016672), the Jaime Erin Follicular Lymphoma Research Consortium (MRG, SN), the Futcher Foundation (LN, MRG), and an MD Anderson Institutional Research Grant (LW). MRG is a Scholar of the Leukemia and Lymphoma Society. HY is a fellow of the Leukemia and Lymphoma Society. PS is supported by a Lymphoma Research Foundation Career Development Award.

