## Supplementary Methods for "Immune microenvironment subtypes and association with tumor cell mutations and antigen expression in follicular lymphoma"

<u>SUPPLEMENTARY FIGURES</u>

Fig. S1: UMAP visualizations of B cells from all patients.

Fig. S2: Unsupervised sub-clustering of T and NK cells from all patients.

Fig. S3: Potential developmental trajectories for CD4 T-cells and CD8 T-cells from all patients inferred by Monocle 3 analysis.

Fig. S4: Independent cohort validation of CD4 and CD8 T-cells subpopulations

Fig. S5: Immunoglobulin hierarchy vs MHCII expression

<u>SUPPLEMENTARY METHODS</u>

Supplementary References

<u>SUPPLEMENTARY TABLES</u>

Table S1: Patient and sample characteristics

Table S2: Cluster marker genes for all cells included in this study

Table S3: Cluster marker genes for CD4 and CD8 T-cell subpopulations

Table S4: DEGs in CD4 CTLs between FL and RLT samples

Table S5: Variants called from whole exome sequencing

Table S6: Differentially expressed genes between mutant samples versus wild-type samples for genes *KMT2D*, *EZH2* and *CREBBP*, respectively.

Table S7: Association between tumor infiltrating T-cell subsets, mutations and MHC status

Table S8: DEGs for CD4 T-cells by MHCII status

Table S9: DEGs for CD8 T-cells by MHCII status

Table S10: Significance levels of genes shown in Figure 4 panel I

### SUPPLEMENTARY METHODS

***Sample collection and single-cell preparation***

Follicular lymphoma tumor specimens were obtained with informed consent in accordance with protocols approved by review board of University of Texas MD Anderson Cancer Center. All FL samples were mechanically dissociated into single-cell suspension. PBS containing 0.04% UltraPure$^{TM}$ Bovine Serum Albumin (BSA, 50mg/ml) was used for cell washing and resuspension to minimize cell losses and aggregation. Cell viability was assessed by trypan blue examination, and samples with more than 80% viable cells were chose for Chromium Single Cell Immune Profiling Solution, according to Chromium Single Cell 5' Library & Gel Bead Kits User Guide (v1 Chemistry). Briefly, the final cell concentration was adjusted to ~1000 cells/µl, single cell suspension mixed with reverse transcription (RT) master mixture were loaded on a 10X Genomics single cell instrument. GEMs were broken and single-strand cDNA was cleaned up with DynaBeads. Amplified cDNA quality and quantity were assessed by High Sensitivity D5000 DNA Screen Tape analysis (Agilent Technologies) and Qubit dsDNA HS Assay Kit (Thermo Fisher Scientific).

**Single-cell RNA-Library construction and sequencing**

We used the 10X Genomics Single-Cell 5' Library Kit (PN-1000020) to construct indexed sequencing libraries following the manufacturer's protocol. Sequencing with indexing the Chromium i7 Samplex index Kit (PN-120262) was conducted on an Illumina Hiseq4000 sequencer with 2×100 bp paired reads to achieve a depth of at least 50,000 read pairs per cell.

**Single-cell V(D)J Enrichment library construction and sequencing**

Chromium Single Cell V(D)J Enrichment Kit (Human B Cell, PN-1000016; Human T Cell, PN-1000005) from 10X Genomics was employed to enrich immune repertoire, T-cell receptor (TCR) or B-cell immunoglobulin (Ig) transcripts. 50 ng of enriched BCR or TCR products was used for library construction, according to manufacturer's protocol. Single Cell V(D)J enriched library was indexed by Chromium i7 Samplex index Kit (PN-120262) and multiple samples were pooled and sequenced in a lane of an Illumina HiSeq 4000 at 2x150bp with a minimum depth of 5,000 read pairs per cell.

**Single-cell targeted genes (CapID) sequencing**

Details were described in our previous publication[1]. Briefly, human immunology signature panel was synthesized by Integrated DNA Technologies (IDT), Inc. Hybridization capture of 5' gene expression (GEX) library was performed using xGen hybridization and wash kit (IDT). The captured library was sequenced on an Illumina HiSeq4000 with 2×100bp paired reads to achieve a depth of at least 50,000 read pairs per cell.

#### Single cell RNA-sequencing bioinformatics

*Raw sequencing data processing, QC, data filtering, and normalization.*

The raw single cell RNA sequencing (scRNA-seq) data were pre-processed (demultiplex cellular barcodes, read alignment (genome reference build: hg19), and generation of feature-barcode matrix) using Cell Ranger (10x Genomics, v3.0.2). Detailed QC metrics were generated and evaluated. Genes detected in <3 cells and cells where < 200 genes had nonzero counts were filtered out and excluded from subsequent analysis. Low quality cells where >15% of the read counts derived from the mitochondrial genome were also discarded. In addition, cells with number of detected genes >6,000 were discarded to remove likely doublet or multiplet captures. The resulting cells were further filtered using the following criteria to clean additional possible doublets: 1) the cells with both productive BCRs and TCRs were removed. 2) For the T, NK, Myeloid and B cells derived from unsupervised clustering analysis, we further cleaned out cells expressing discrepant canonical markers. For example, in the T and NK cell lineages, the cells or cell clusters that have productive BCRs, or express lineage-specific B or Myeloid cell markers were removed. Similarly, in the Myeloid cell populations, the cells or cell clusters that have productive TCRs/BCRs, or expressing canonical T or B cell markers were removed. Finally, cell clusters that have productive TCRs, or express T or Myeloid cell lineage markers were removed from malignant/non-malignant B cell lineages. To reduce the batch effects, we performed batch effect correction on each cell population of non-malignant cells (including CD4 T, CD8 T, proliferating T, NK, Myeloid and non-malignant B cell clusters) with Harmony[2]. The results of principal component analysis (PCA), Uniform Manifold Approximation and Projection (UMAP)[3] [arXiv preprint arXiv:1802.03426] plots and sample-by-cluster distribution were carefully reviewed. Seurat[4] version 3 was applied to the filtered gene-cell matrix to generate the normalized UMI counts using *NormalizeData* function.

*Unsupervised cell clustering and dimensionality reduction.*

Seurat version 3 was applied to the normalized gene-cell matrix to identify highly variable genes. The elbow plot was generated with the *ElbowPlot* function of Seurat[5] and based on which, the number of significant principal components (PCs) were determined. Different resolution parameters for unsupervised clustering were then examined in order to determine the optimal number of clusters. For this study, the first 100 PCs calculated using the 2000 most highly variable genes identified by Seurat were used for unsupervised clustering analysis with the resolution set to 0.5, yielding a total of 31 cell clusters. Dimensionality reduction and 2-D visualization of the single cell clusters was performed using UMAP[3] with Seurat function *RunUMAP*. In our data, the batch effect is not obvious as non-malignant cells from different samples were observed to be clustered together. For non-malignant cell populations, the batch effect correction was performed using Harmony[2].

*Determination of major cell types and cell states.*

To determine the cell types and cell states, cluster marker genes were calculated using *FindAllMarkers* function in Seurat R package. The significant DEGs (FDR q-value < 0.05, Fold change > 1.2) were examined and an integrative approach was used to determine cell types and states. The major cell type (CD4 and CD8) was defined by marker gene expression (*CD3D*, *CD4*, *CD40LG*, *CD8A*, *CD8B*) by 10X transcriptome and CapID sequencing data. The functional state of each single cell (activated, memory, exhausted, regulatory) was determined using markers described by Sade-Feldman *et al*[6] and Chunhong Zheng *et al*[7]. To describe the cell types and states that were defined by each cluster, we performed a manual review of the differentially expressed genes (DEGs) that were identified for each cell cluster by Seurat.

*Inferring cell cycle stage, hierarchical clustering, differentially expressed genes (DEGs).*

The cell cycle stage was computationally assigned to each individual cell using the R code implemented in Seurat based on expression profiles of the cell cycle-related signature genes, as previously described[8]. DEGs were identified for each cluster using the *FindMarkers* function of in Seurat R package and DEG list was filtered with the default criteria: the gene should be expressed in 10% or more cells in the more abundant group; the absolute expression fold change >1.2; and FDR q-value <0.05. Hierarchical clustering was performed for each cell type using the Ward's minimum variance method. Heat map was then generated using the *heatmap* function in pheatmap R package for filtered DEGs.

*TCR V(D)J sequence assembly, paired clonotype calling, TCR diversity and clonality analysis and integration with scRNA-seq data.*

Cell Ranger v3.0.2 for V(D)J sequence assembly was applied for TCR reconstruction and paired TCR clonotype calling. The CDR3 motif was located and the productivity was determined for each single cell. The clonotype landscape was then assessed and the clonal fraction of each identified clonotype was calculated. The TCR clonotype diversity matrix was calculated using the tcR R package[9]. TCR clonality was defined as 1-Peilou's evenness and was calculated on productive rearrangements as previously described[10]. Clonality values approaching 0 indicate a very even distribution of clone frequencies, whereas values approaching 1 indicate an increasingly asymmetric distribution in which a few clones are present at high frequencies. The TCR clonotype data was then integrated with the T-cell phenotype data inferred from single cell gene expression analysis based on their shared unique cell barcodes.

*Analysis of large-scale copy number variations.*

To quantify the level of aneuploidy, profiles of copy number variation (CNV) generated by inferCNV (https://github.com/broadinstitute/inferCNV) were aggregated using a similar strategy adopted by a previous study[11]. We first computed arm-level CNV scores as the mean of the squares of CNV values across each chromosomal arm. The arm-level CNV scores were further aggregated across all chromosomal arms by taking the average of the arm-level scores.

***Whole-exome sequencing (WES) data processing and genotyping quality check (QC).***

*Library preparation and hybrid capture sequencing.*

Genomic DNA (gDNA) was extracted from these remaining cells of prepared 10X single cell suspension using AllPrep DNA/RNA Mini Kit (QIAGEN). 50–200ng of gDNA were applied to DNA library preparation using KAPA HyperPlus Kit (Roche), according to the manufacturer's protocol. TruSeq adapters (Bioo Scientific) were utilized at the recommended ratio to input DNA. Library quality was assessed by High Sensitivity D1000 DNA Screen Tape analysis (Agilent Technologies) and Qubit dsDNA HS Assay Kit (Thermo Fisher Scientific). Libraries were 6-plexed in equal quantities and a 1µg of pooled libraries were enriched by hybrid capture using a Nimblegen SeqCap Exome v3 (Roche), according to the manufacturer's protocol. Each capture pool was sequenced on a lane of a HiSeq4000 to generate 2 x 100bp reads.

*Somatic mutation calling, filtering, and functional annotation.*

Mutation detection and filtering was performed as previously described[12]. Raw FASTQ files were assessed for quality using FASTQC. Samples with high quality metrics were run through our in-house pipeline. FASTQ files were (i) aligned to the human genome (hg19) using BWA-Mem[13], (ii) deduplicated using Picard MarkDuplicates, (iii) realigned around InDels using GATK[14], and (iv) recalibrated by base score using GATK[14]. On-target rate and coverage over the targeted region were calculated by Picard CalculateHSMetrics. Only samples achieving >50X average on-target coverage were utilized. For samples with additional sequencing performed to obtain sufficient coverage, FASTQC was performed on individual sequencing events and bam files from the same sample were merged using BamTools Mappings Merger following alignment. Variants were called by GATK Unified Genotyper and VarScan2[15]. Only variants called by both tools, with a minimum coverage of 30X and ≥3 supporting reads were retained, which we have previously shown to provide a sensitivity of 96.7% and specificity of 92.9%[16]. All variants were annotated using SeattleSeq[17]. To avoid mapping artifacts in repetitive regions, all variants within RepeatMasker or tandemRepeat annotated regions were filtered from the dataset. All variants in dbSNP build 32 or the Genome Aggregation Database (gnomAD) were removed[18] to control for potential germline variants. Only genes with mutations present within ≥3 tumors were considered for analysis.

##### *Public datasets*

Gene signatures were validated on 15 bulk FL cohorts: GSE127462, GSE53820, GSE93261, ICGC_MALY_DE, GSE66166, GSE55267, GSE37088, E-MTAB-6088, E-MEXP-2305, GSE132929, GSE103944, GSE32018, GSE48047, GSE142334 and GSE16131. RNA-seq datasets were processed using Kallisto[19]. Gene signatures were quanitified using ssGSEA[20]. Non-FL samples from these cohorts (if present) were removed before ssGSEA analysis. Gene signature scores were median scaled within each dataset and combined to increase the number of samples (n=1,269).

To validate the identified signatures, we used a public scRNA-seq dataset including B-cell lymphoma data[21]. The data were downloaded and analyzed according to best practice[22]. In brief, cells with high mitochondrial expression (> 10% of total UMIs per cell) were excluded from further analysis. A subset of T-cells (n = 8,589) was selected using the provided annotation and re-analyzed. Immunoglobulin and T-cell receptor gene expression data were excluded from further analysis. After the selection of 2,000 highly variable genes and the removal of expression linked with the total UMI counts and the percent of mitochondrial UMIs, cells from different patients were integrated using Harmony[2]. Using the Leiden algorithm[23] (with resolution = 1.5, based on the community graph constructed on first 15 Harmony components and with 50 local neighborhoods). Gene signature scores were calculated using the scanpy[24] package function *score_genes*.

##### *Statistical analysis*

In addition to the bioinformatics approaches described above for scRNA/TCR-seq data analysis, all other statistical analysis was performed using statistical software R v3.5.2. Analysis of differences in immunological features (continuous variables) between patient groups (MHC II high vs. MHC II low) was determined by the nonparametric Mann-Whitney U test. To control for multiple hypothesis testing, we applied the Benjamini-Hochberg method to correct p values and the false discovery rates (q-values) were calculated. All statistical significance testing was two-sided and results were considered statistically significant at p-value < 0.05.

#### SUPPLEMENTARY FIGURES

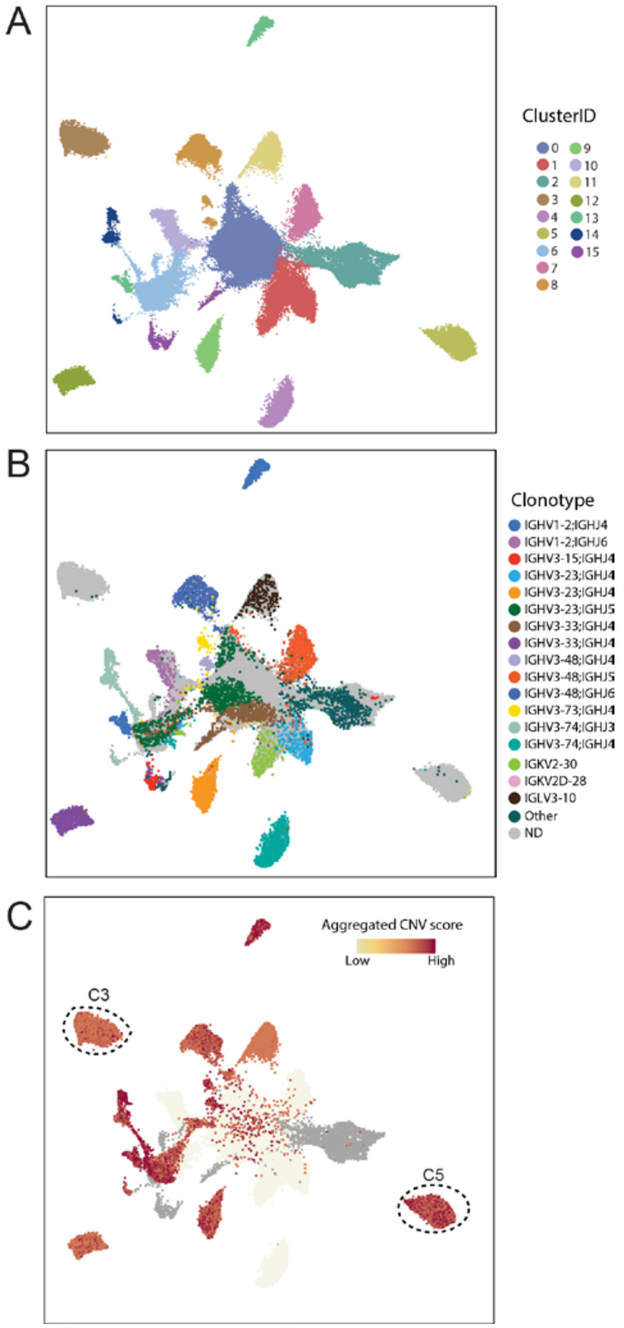


**Supplementary Fig. S1. UMAP visualizations of B cells from all patients.** The UMAP visualization of B cells are colored by cluster ID (A), the clonotypes defined with VDJ sequencing (B), and the inferred copy number variation (CNV) scores (C).

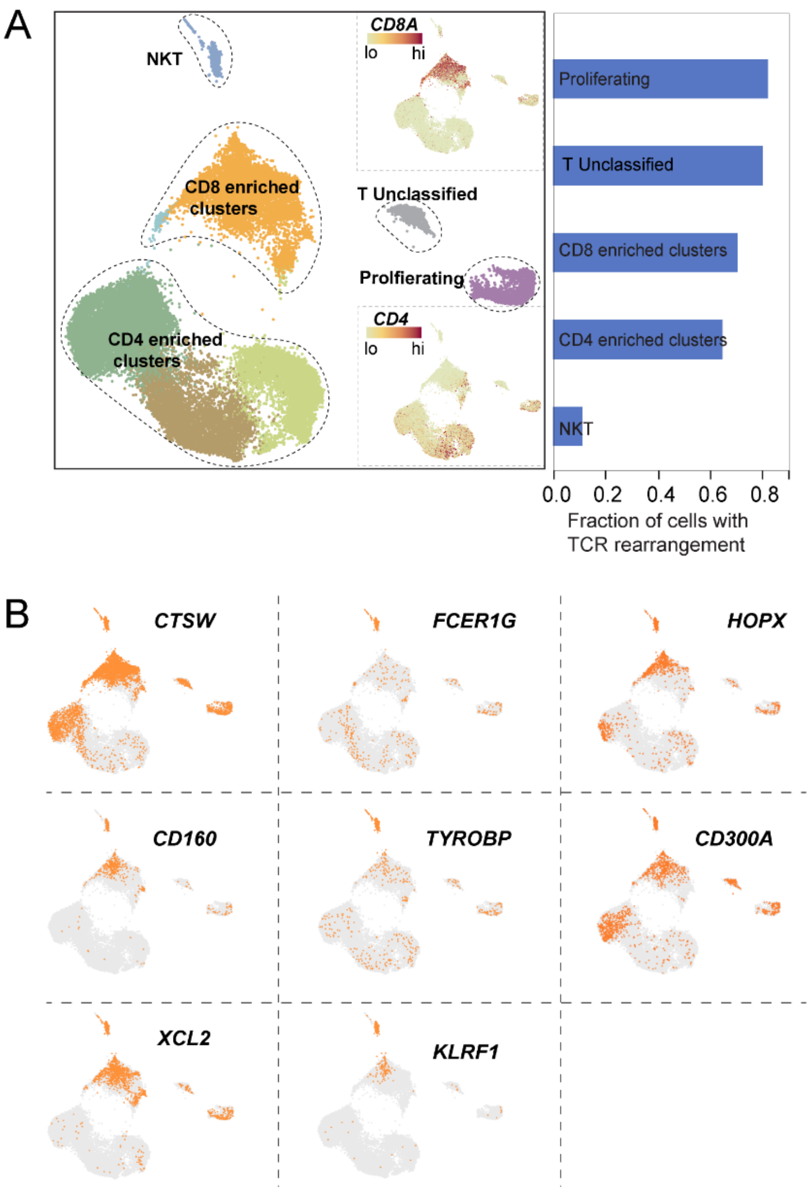


**Supplementary Fig. S2. Unsupervised sub-clustering of T and NK cells from all patients.** **A,** UMAP visualization of T cells and NK cells from all patients colored by cluster ID (left) and the barplot showing fraction of cells with detectable TCR rearrangement per cell type based on scTCR-seq (right). Feature plots of CD8A and CD4 are shown in insets at top right and bottom right, respectively. Lo, low expression; hi, high expression. **B,** Feature plots of representative genes. Orange colored cells denotes the corresponding gene expression level (logUMI) > 0.

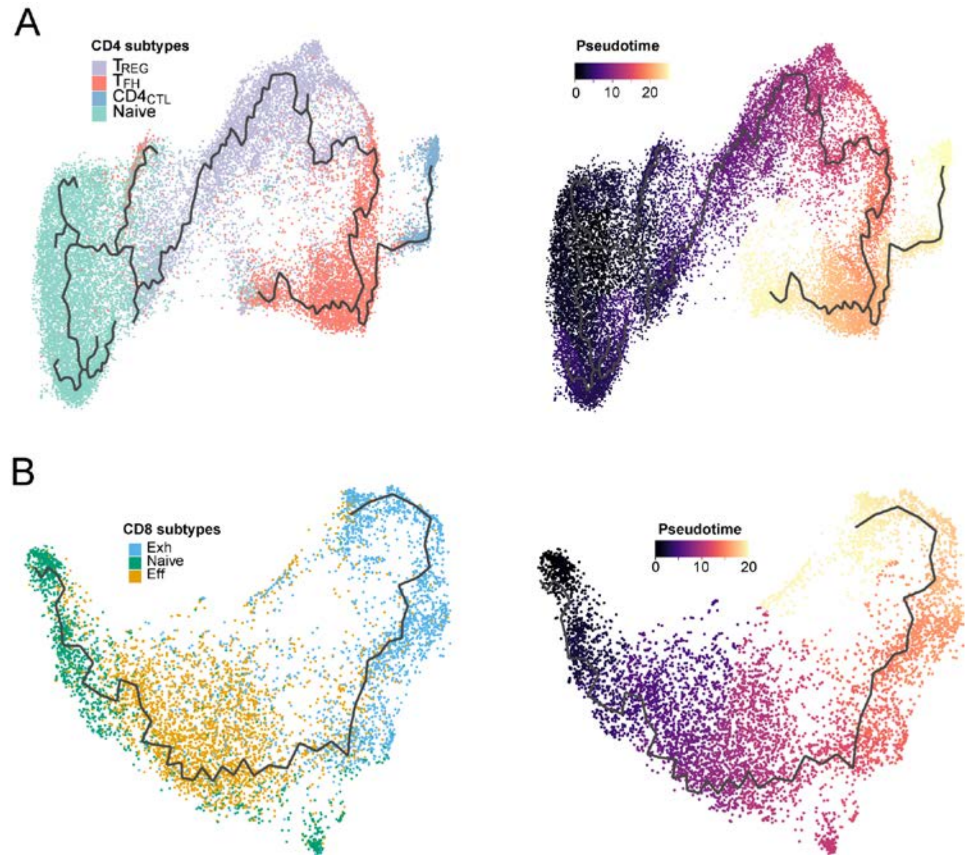


**Supplementary Fig. S3. Potential developmental trajectories for CD4 T-cells and CD8 T-cells from all patients inferred by Monocle 3 analysis. A,** Monocle trajectory plots showing cells colored by CD4 T-cell subpopulations (left) and by inferred pseudotime based on global expression profiles (right). **B**, Monocle trajectory plots showing cells colored by CD8 T-cell subpopulations (left) and by inferred pseudotime (right).

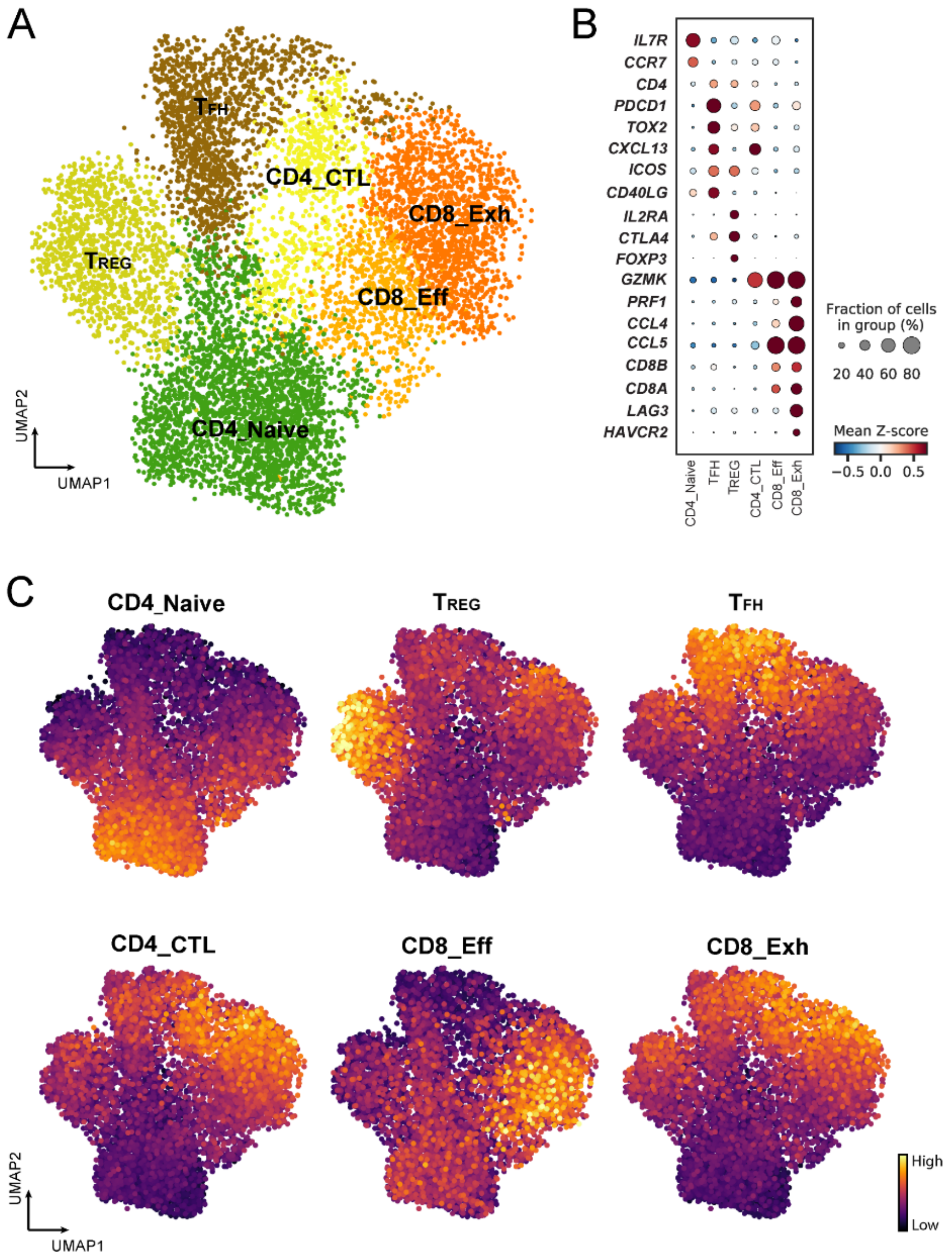


**Supplementary Fig. S4. Independent cohort validation of CD4 and CD8 T cells subpopulations. A,** UMAP visualization of the subpopulations identified for CD4 and CD8 T-cells from the independent cohort from Roider et al.[21]. Cells are colored by their assigned cell types. The number of cells in this validation cohort is 8,453 cells. **B**, Bubble plot showing expression of marker genes. Both the fraction of cells expressing signature genes (indicated by the size of the circle) as well as their scaled expression levels (indicated by the color of the circle) are shown. **C**, Feature plots showing the cell subpopulation signature scores calculated using DEGs obtained in this study.

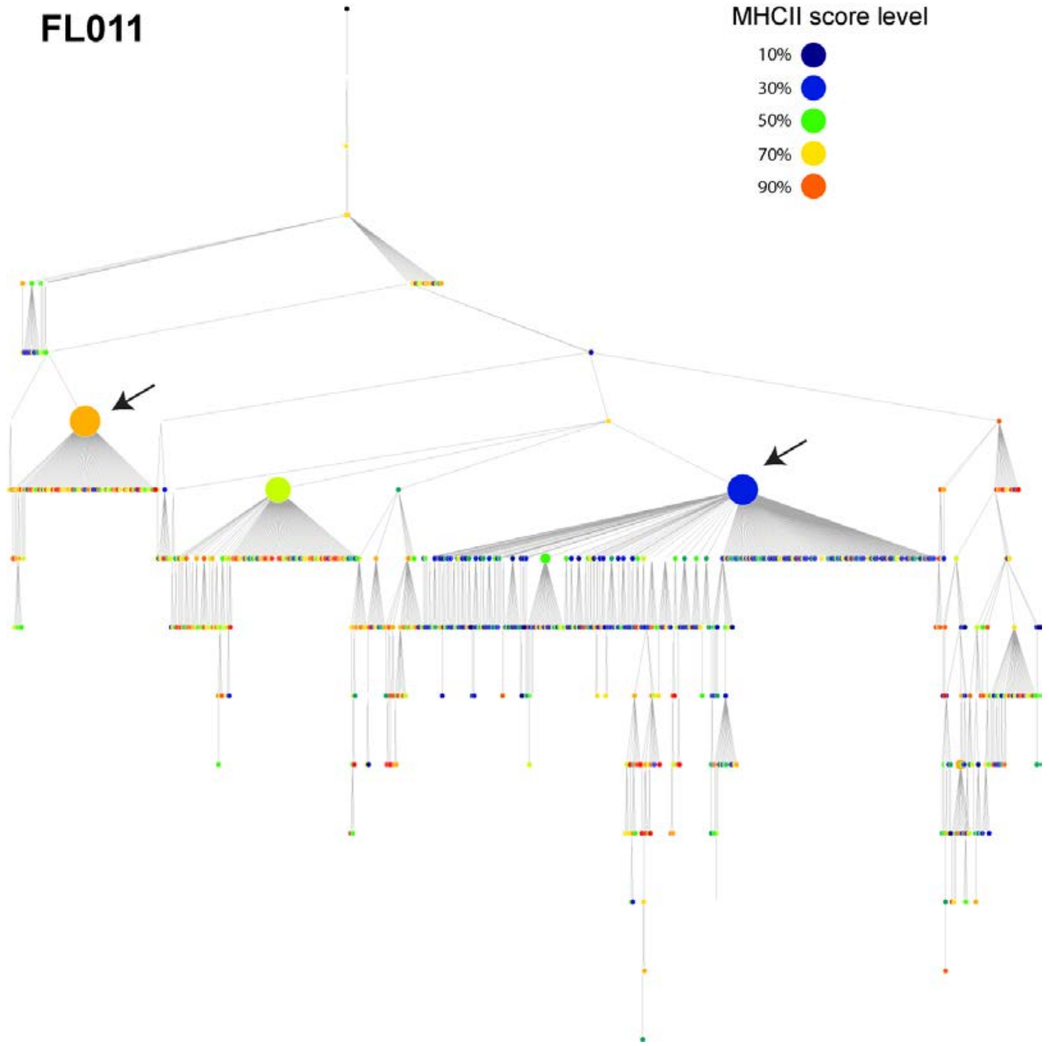


**Supplementary Fig. S5. Correlation between MHCII expression scores and the immunoglobin hierarchy.** The immunoglobulin hierarchy tree constructed of one patient (ID: 044) is correlated with the MHCII scores defined using the differentially expressed MHC II genes identified in this study.
